# Eco-evolutionary control of pathogens

**DOI:** 10.1101/858621

**Authors:** Michael Lässig, Ville Mustonen

## Abstract

Control can alter the eco-evolutionary dynamics of a target pathogen in two ways, by changing its population size and by directed evolution of new functions. Here we develop a fitness model of eco-evolutionary control that specifies a minimum leverage for successful control against the intrinsic dynamics of the pathogen. We apply this model to pathogen control by molecular antibody-antigen binding with a tunable level of antibodies. By analytical solution, we obtain a phase diagram of optimal control and show that an error threshold separates regimes of successful and futile control. Our analysis identifies few, independently measurable fitness parameters that predict the outcome of control. We show that optimal control strategies depend on mutation rate and population size of the pathogen, and we discuss how monitoring and computational forecasting affect the efficiency of control. We argue that these results carry over to more general systems and are elements of an emerging eco-evolutionary control theory.

## Introduction

Control of human pathogens is a central goal of medicine. Important examples are antimicrobial and antiviral therapies and vaccinations; similarly, cancer therapies aim to control tumor cell populations. Biological hosts, notably the human immune system, face related issues of pathogen control. In most cases, control targets pathogen populations with fast-paced replication and evolution. Its goal is to alter these dynamics: to prevent or elicit an evolutionary process of the pathogen, or to curb the pathogen population by reducing its ecological niche. Pathogen control has seen spectacular successes, e.g., in the eradication of smallpox and in HIV combination therapies [1]. But control is often compromised by escape evolution of the pathogen, highlighting the importance to factor pathogen evolution into control protocols [2, 3]. Promising evolutionary avenues include adaptive pathogen control and cancer therapy [4, 5, 6], vaccination, drug development and immunotherapy strategies based on evolutionary predictions [7, 8, 9, 10], and controlled evolution of immune antibodies [11, 12, 13]. However, we lack quantitative relations between leverage and cost of control to generate optimization criteria and protocols that are comparable across systems. These are central elements of an eco-evolutionary control theory.

Because population dynamics and evolution are stochastic processes, any eco-evolutionary control operates on the likelihood of future states. Successful control turns a likely process into an unlikely one (e.g., the evolution of antibiotic resistance) or vice versa (e.g., the evolution of a broadly neutralizing antibody). In a broader scientific context, directing a stochastic process towards a future objective is a classic subject of control theory [14, 15]. There is a well-established conceptual and computational framework to optimize control protocols, given complete knowledge of the dynamical rules and the ability to forecast likely future outcomes. However, the swords of eco-evolutionary control are blunter, and establishing an appropriate control theory faces new challenges. First, the control of an evolving population is based, at best, on limited dynamical information and forecasting capabilities. For human interventions, optimizing control is inextricably linked to predictive evolutionary analysis, which is a topic of high current interest but far from a comprehensive understanding [16]. Second, control theory has to factor in the underlying biological mechanism of control. Host-pathogen interactions are often based on bio-molecular interactions, such as drug-target or antibody-antigen binding [17]. This imposes specific constraints on control forces and their leverage on the pathogen system, which are discussed below. Third, developing an appropriate dynamical model of control update and pathogen response calls for a merger of control theory with ecological dynamics and population genetics. These questions are the topic of the present paper.

In the first part, we focus on pathogen dynamics under control. The driving forces of these dynamics are the pathogen’s intrinsic fitness and entropic factors (such as the mutational target of a pathogen trait), as well as the additional, host-dependent selection generated by control. We derive general minimum-leverage relations that specify the strength of control needed to alter the evolution of the pathogen towards the host’s control objective. In the second part, we add the control dynamics of the host, which is governed by its cost-benefit balance. We analyze the coupled host-pathogen dynamics in a minimal model of bio-molecular control, in which the host produces antibodies that bind to the pathogen. This model captures two complementary control modes, which are associated with different host-pathogen interactions. For ecological control, bound antibodies impede pathogen growth. The control objective is to reduce the pathogen’s carrying capacity, a deleterious collateral is the evolution of resistance. For evolutionary control, bound antibodies reduce pathogenicity. The control objective is the adaptive evolution of an antibody binding site (epitope) in the pathogen population, a collateral is the concurrent increase of the carrying capacity.

From an analytical solution of this model, we obtain efficiency phase diagrams that map parameter regimes of efficient and futile control. Their most pronounced feature is an *error threshold*, which marks a switch of the pathogen’s evolutionary state and a rapid change of control efficiency. The error threshold depends jointly on the host’s control mechanism and the pathogen’s mutation rate and population size, highlighting the link of control and population genetics. We discuss implications of these results for biomedical applications: how information processing impacts mode and efficiency of control, and how measurements of core pathogen and host data can be used to predict control outcomes.

## Eco-evolutionary control theory

### Eco-evolutionary dynamics

We consider a population of pathogens with a quantitative trait *G*, which has a peaked trait distribution *ρ*(*G*) characterized by its mean Γ and variance Δ. We describe the evolution of the trait and the concurrent population dynamics in an ecological niche by coupled stochastic equations for the mean Γ and the population size *N*,

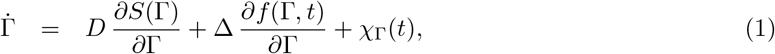

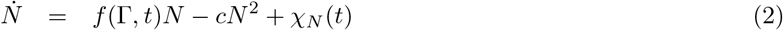

with white noise *χ*_Γ_(*t*), *χ*_*N*_ (*t*) of mean and variance

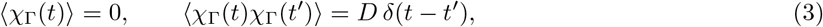

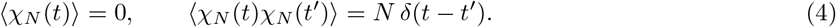

The trait dynamics of Eq. (1) depend on the entropy landscape *S*(Γ), which is defined as the log density of states with trait value Γ, and the fitness seascape *f*(Γ, *t*), which is often explicitly time-dependent. The trait diffusion constant, *D* = *Uϵ*^2^, is set by the total rate *U* and the mean square trait effect *ϵ*^2^ of mutations at genomic loci encoding the trait [18, 19]. The trait response to selection, Δ = 2*DN*_*e*_, is also proportional to the effective population size *N_e_*, which equals the coalescence time of the evolutionary process [19, 20]. This quantity depends on the population size dynamics in a model-dependent way, often generating an inhomogeneous response to selection, Δ(Γ) = 2*DN*_*e*_(Γ). The population dynamics of Eq. (2) depend on the fitness *f*(Γ, *t*), which sets the basic reproductive rate, and the constraint parameter c of the ecological niche. Given a static or slowly varying fitness function, these dynamics generate population size fluctuations around a carrying capacity 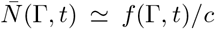. Details of this stochastic calculus are given in Methods.

Recording the eco-evolutionary process in an individual population over a time interval (*t_i_*, *t_f_*) generates a trait path **Γ**, which is given by a time-dependent trait Γ(*t*) with initial value Γ_*i*_ and final value Γ_*f*_. We define the evolutionary flux

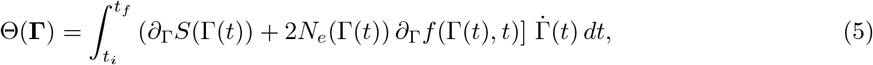

which sums the entropy increments dS and the scaled fitness increments 2*N*_*e*_*df* generated by trait changes along the path **Γ**. The flux Θ becomes an important building block of eco-evolutionary control theory: it characterizes the likelihood of an evolutionary path and its alteration by control. This is shown in Methods, building on previous results in evolutionary statistics and stochastic thermodynamics [21, 22]. Pathogen evolution often involves large flux amplitudes (Θ ≫ 1), which can be generated by strong selection or sufficiently complex evolutionary traits. In this case, we obtain a deterministic criterion: observable evolutionary paths have a positive flux,

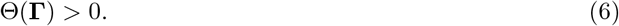

This holds not only for the full process, but for any sub-period 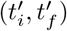 (as long as flux amplitudes remain large).

### Minimum leverage relations

Eco-evolutionary control is exerted by altering selection: the controlled system is governed by a fitness seascape *f*(Γ, *t*) = *f*_*b*_(Γ) + *f*_*c*_(Γ, *t*), which contains the background component *f*_*b*_(Γ) and the control component *f*_*c*_(Γ, *t*). We now use the flux inequality (6) to determine lower bounds on *f*_*c*_(Γ, *t*), i.e., a minimum leverage of the controlling onto the controlled system required for successful control.

First, a stationary control landscape *f*_*c*_(Γ) can *maintain* a controlled trait value Γ* against a reference value Γ_0_ if the flux condition (6) is fulfilled for any evolutionary path **Γ** from Γ_0_ to Γ*. Writing the flux as the differential of a free fitness landscape, Θ(Γ) = *ψ*(Γ_*f*_) − *ψ*(Γ_*i*_), we obtain the minimum-leverage relation for maintenance of a trait,

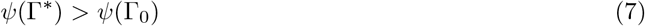

(Methods). This condition says that escape mutations from Γ* to Γ_0_ are suppressed by negative selection. It applies to large populations where such escape mutations would otherwise be frequent, but it can be undercut in small populations where escape mutations are rare (see below). In the case of a constant *N_e_*, the inequality (7) takes the simpler form

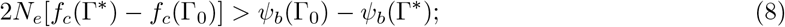

i.e., the scaled control fitness leverage has to exceed the drop in free fitness of the uncontrolled system, which is the sum of the scaled fitness difference *f_b_*(Γ_0_) − *f_b_*(Γ*) and the entropy difference *S*(Γ_0_) − *S*(Γ*) (Methods). In Fig. 1, we illustrate the minimum-leverage relation (8) for two control scenarios: ecological control aimed at reducing 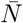 while maintaining the wild type trait Γ_wt_ against the evolution of a resistant state Γ_*e*_ (Fig. 1A), and evolutionary control aimed at maintaining an evolved equilibrium Γ_*e*_ against reversal to Γ_wt_ (Fig. 1B). In these examples, the evolution of a resistance trait under ecological control is a loss of function, which often implies *S*(Γ_*e*_) > *S*(Γ_wt_), whereas evolutionary control elicits a gain of function with *S*(Γ_*e*_) < *S*(Γ_wt_). Below, we will use the condition (8) to map strong and weak control regimes.

Second, a time-dependent control seascape *f_c_*(Γ, *t*) can *elicit* a trait value Γ* from a wild-type value Γ_0_, if there is a path **Γ** from Γ_0_ to Γ* that fulfils the flux condition (6) for any segment covered in an arbitrary sub-period 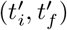. This leads to the local minimum-leverage condition for evolutionary control,

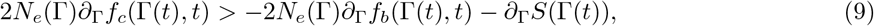

which requires uphill movement of the trait Γ(*t*) in the local gradient 2*N_e_*(Γ)*df* (Γ, *t*)+*dS*(Γ). In particular, the control must bridge fitness valleys of the background landscape *f_b_*(Γ), which often requires high transient control amplitudes. The deterministic minimum leverage condition (7) is valid up to small fitness troughs that can be crossed by fluctuations at low effective population size.

**Figure 1:**
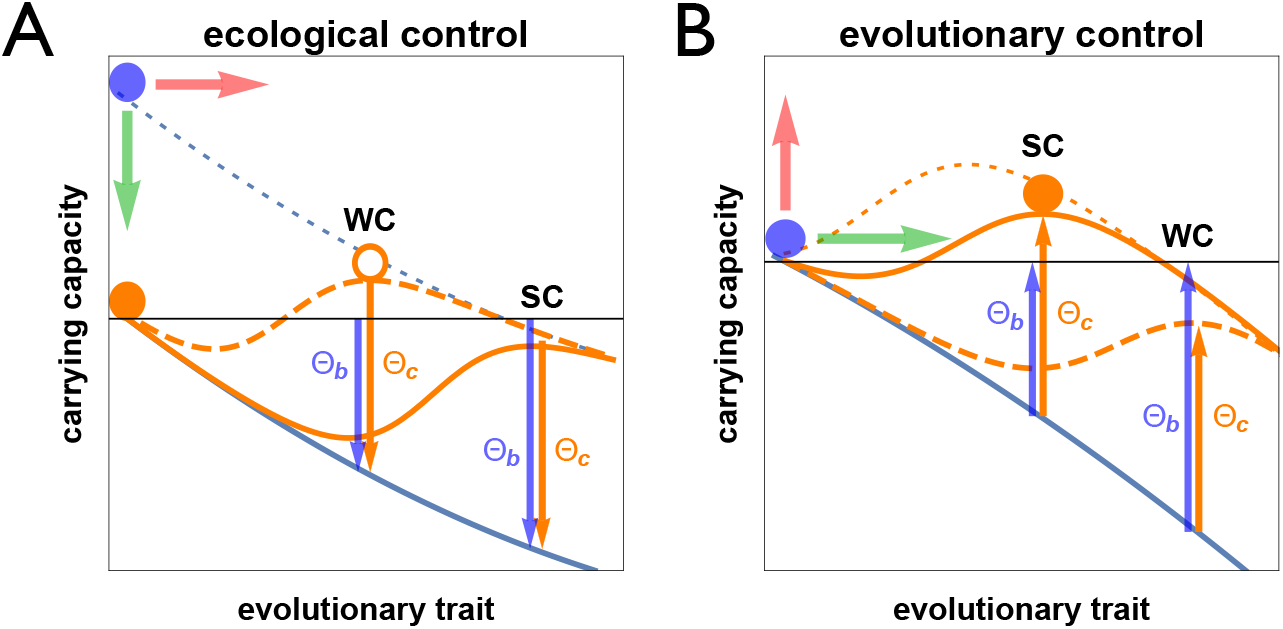
Modes and leverage of eco-evolutionary control. A pathogen with population mean trait Γ under control with amplitude *ζ* lives in a free fitness landscape *f* (Γ, *ζ*) (orange lines), which is the sum of a background component *ψ*_*b*_(Γ) (blue line) and a control landscape *f_c_*(Γ, *ζ*). (A) Ecological control, starting from an uncontrolled wild type pathogen (blue dot), has the objective of reducing the pathogen’s carrying capacity (green arrow) – here by antibody binding – and the collateral effect of resistance evolution (red arrow). The effi ciency of control depends on the leverage Θ_*c*_ = 2*N_e_*[*f_c_*(Γ_wt_) − *f_c_*(Γ_*e*_)] (orange arrows), which is equal in magnitude to the fitness effect of antibody binding on the pathogen, and on the background effect of resistance, Θ_*b*_ = *ψ*_*b*_(Γ_wt_) − *ψ_b_*(Γ_*e*_) (blue arrows), which contains the fitness cost of the resistance trait and the entropy difference between the wild type trait Γ_wt_ and the optimal evolved trait Γ_*e*_. Strong control (SC, Θ_*c*_ > Θ_*b*_, solid orange line) suppresses the evolution of resistance and constrains the pathogen to the wild type (orange dot); weak control (WC, Θ_*c*_ < Θ_*b*_, dashed orange line) triggers the evolution of resistance (orange circle). Downward arrows represent negative numbers. (B) Evolutionary control has the objective of eliciting a new pathogen trait (green arrow) and the collateral of increasing its carrying capacity (Γed arrow). Dynamical control elicits the evolved trait along a path of positively selected trait increments, which requires elevated transient control amplitudes (dotted orange line). Stationary control aims at maintaining the evolved trait Γ_*e*_; its effi ciency depends on the leverage Θ_*c*_ = 2*N_e_*[*f_c_*(Γ_*e*_) − *f_c_*(Γ_wt_)] (orange arrows) and on the background effect Θ_*b*_ = *ψ*_*b*_(Γ_*e*_) − *ψ*_*b*_(Γ_wt_) (blue arrows). Strong control (SC, Θ_*c*_ > Θ_*b*_, solid orange line) maintains the evolved trait (orange dot); weak control (WC, Θ_*c*_ < Θ_*b*_, dashed orange) triggers a reversal to the wild type.

The inequalities (7) – (9) specify the minimum leverage that a controlling host system must exert on the controlled pathogen system, in order to elicit a feature that would not evolve spontaneously and to maintain this feature against reverse evolution towards the wild type. These relations are formally related to the maximum-work theorem of thermodynamics, which specifies the minimum work uptake (or maximum work release) associated with a given free energy change of a thermodynamic system. Unlike in thermodynamics, however, the minimum-leverage relations say nothing about cost and benefit of control for the the controlling system. This requires a mechanistic model of the host-pathogen control interaction, to which we now turn.

## Application to pathogen control

### Antibody-antigen interactions

We now focus on a specific control scenario, in which a host exerts control by producing antibodies that bind to a pathogen (also referred to as antigen in this context). The probability that the pathogen is bound,

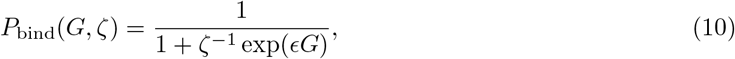

depends on the pathogen trait *G* (with wild type value *G*_wt_ = 0) and the antibody density *ζ* (measured here in units of the dissociation constant or IC50 of the wild-type pathogen). We consider two cases, which cover the control modes illustrated in Fig. 1. For ecological control (*ϵ* = +1), *G* is a resistance trait (the log of the dissociation constant); i.e., a population with evolved resistance (Γ_*e*_ > 0) has reduced binding. For evolutionary control (*ϵ* = −1), *G* is the epitope affinity (the log of the association constant); this mode is to elicit an evolved population (Γ_*e*_ > 0) with increased binding.

### Pathogen and host fitness landscapes

We assume host and pathogen live in coupled fitness land-scapes of the form

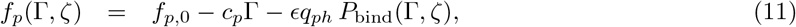

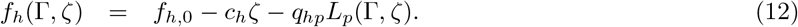

The pathogen has a background fitness *f*_*p*,0_ in the uncontrolled wild type, a background cost *c_p_* > 0 per unit of the trait (the linear form is taken for simplicity), and a binding-dependent control term *f_c_* = −*ϵq*_*ph*_ *P*_bind_ with a selection coefficient *q_ph_* > 0. The host has a background fitness *f*_*h*,0_ in the absence of control and pathogens, a production cost *c_h_* per unit of antibody, and an interaction term depending on the pathogen load *L_p_*(Γ, *ζ*) with a selection coefficient *q_hp_* > 0. In the case of ecological control, the load is generated by the full pathogen population, *L_p_*(Γ, *ζ*) = *f_p_*(Γ, *ζ*), where the pathogen is assumed to be at its carrying capacity 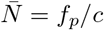 given by Eq. (2) and measured in units of *c*^−1^. In the case of evolutionary control, we use a load function *L_p_*(Γ, *ζ*) = (1 − *P*_bind_(Γ, *ζ*))*f_p_*(Γ, *ζ*), assuming that bound pathogens lose their pathogenicity. Fig. 2 shows the fitness landscapes of Eqs. (11) and (12) for ecological and evolutionary control. In both cases, the pathogen has two local fitness maxima (solid and dashed line) with a rank order depending on the control amplitude.

**Figure 2:**
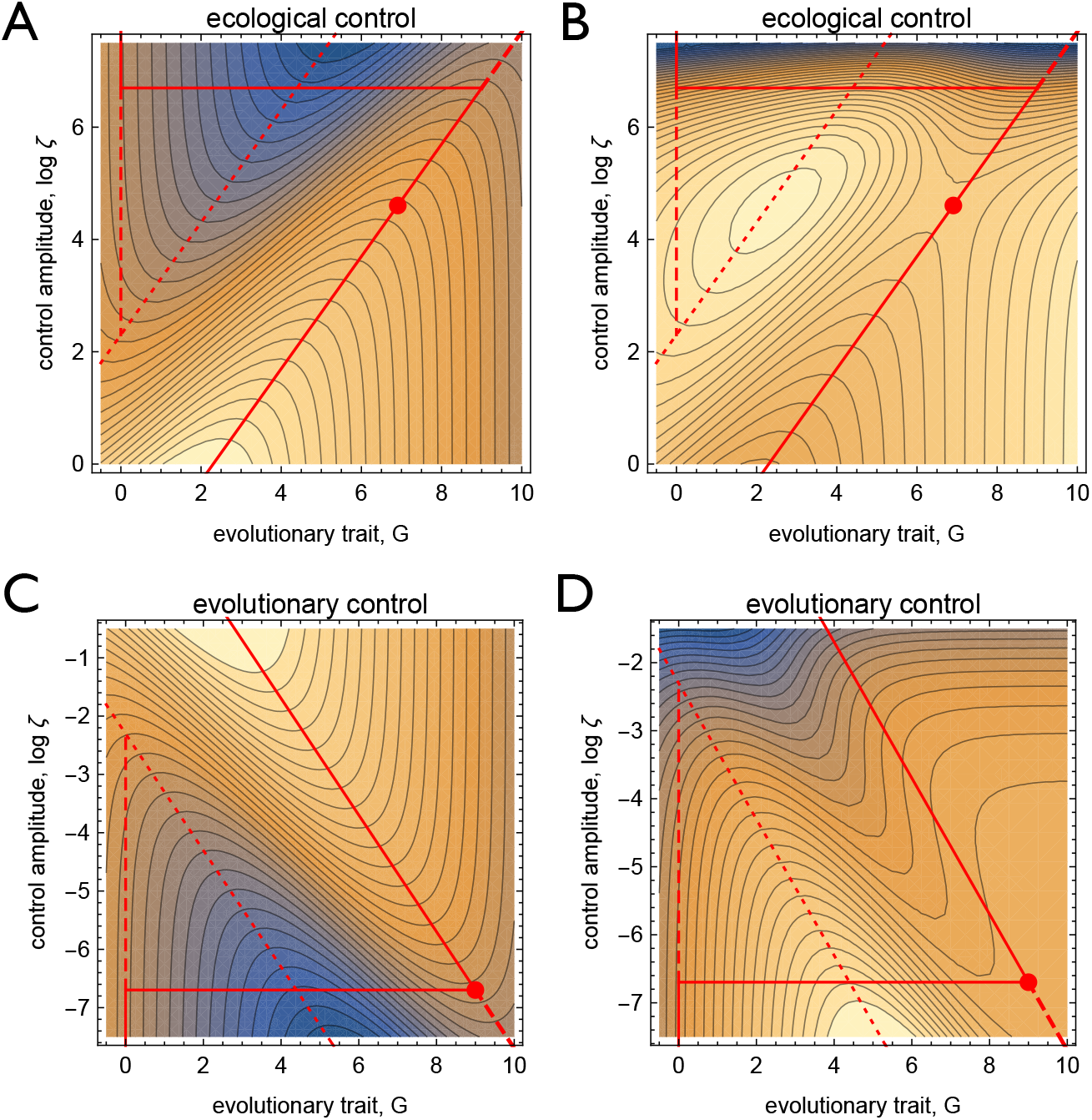
Fitness landscapes of pathogen control. (A) Ecological control, (B) evolutionary control. Pathogen fitness *f_p_* (left) and host fitness *f_h_* (right) are shown as functions of the log control amplitude, log *ζ*, and the pathogen trait, *G*. Specific control loci: pathogen fitness minimum (dotted), pathogen fitness maxima (stable: solid, metastable: dashed, stability switch: horizontal line), optimal control (log *ζ*\*, *G*\*) (dot). Model parameters: *f*_*p*,0_ = 1, *q_ph_* = 1/*q_ph_* = 0.9, *c_p_* = 0.09, *c_h_* = 0.001 (ecological control); *f*_*p*,0_ = 1, *q_ph_* = 1/*q_hp_* = 9, *c_p_* = 0.9, *c_h_* = 1 (evolutionary control).

### Stationary control

We now focus on pathogens that can be contained by long-term control but cannot be killed by a short-time protocol (this is, currently, an appropriate assumption for HIV and some cancers). We first consider the ecological control of a large pathogen population by a stationary landscape *f_c_*(Γ, *ζ*). At a given control amplitude *ζ*, such populations evolve to a conditional equilibrium point Γ*(*ζ*) = arg max_Γ_ *f_p_*(Γ, *ζ*), which maximizes the the pathogen (free) fitness (with a subleading contribution of the entropy *S*(Γ, *ζ*)). The optimal stationary control is then realized at the point

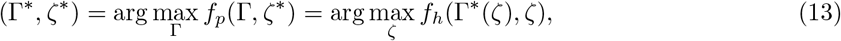

which satisfies the minimum-leverage condition (7) (i.e., is stable against pathogen escape mutations) and maximizes the host fitness. In the fitness landscapes of Eqs. (11) and (12), this control optimization problem can be solved analytically (Methods). The solution reveals the existence of two regimes,

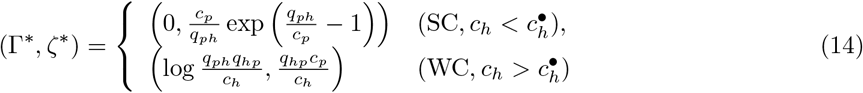

which are separated by an error threshold of the control cost,

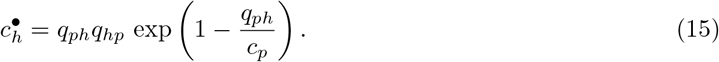

For 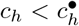, the optimal protocol is in a *strong-control* (SC) regime, which keeps the pathogen in its wild type, i.e., suppresses the evolution of resistance. For 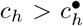, the optimal protocol is in a *weak-control* (WC) regime with evolved pathogen resistance (this case is shown in Fig. 2A). We define the control efficiency

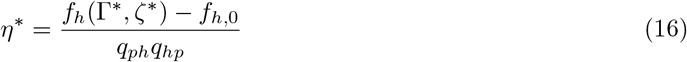

as the gain in host fitness due to control, relative to the maximal gain in the absence of any control cost. Fig. 3A shows the efficiency as a function of the cost parameters (*c_p_*, *c_h_*). The error threshold 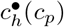 (yellow line) marks a rapid decline of the efficiency from values *η*\* ~ 1 in the SC regime to *η*\* ≪ 1 in the WC regime. At the transition, the resistance trait switches from Γ* = 0 (SC) to Γ_thr_ = *q_ph_*/*c_p_* − 1 (WC), while *f_h_*, *f_p_*, and *ζ*\* remain continuous (Fig. S1).

For stationary evolutionary control, we find a similar emergence of two control regimes with an error threshold

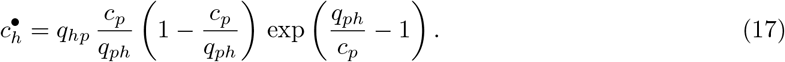

**Figure 3:**
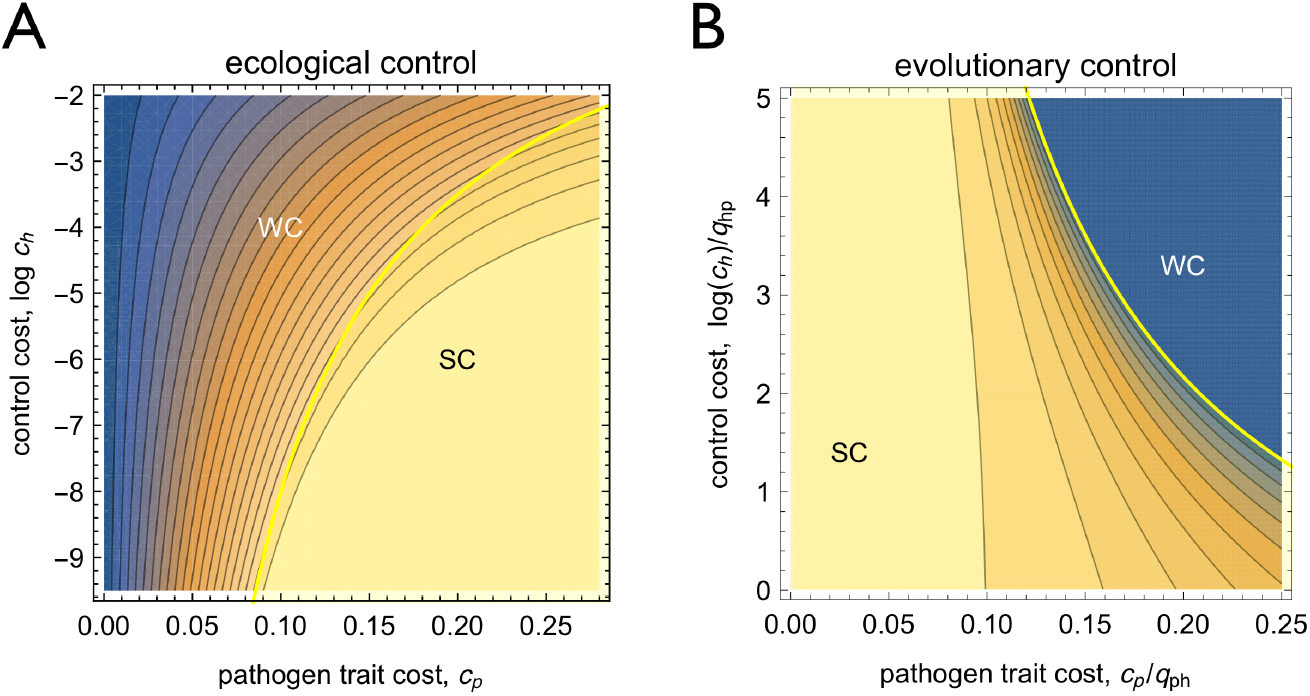
Phase diagrams of stationary control. (A) Ecological control, (B) evolutionary control. The control efficiency *η*\* is shown as a function of the cost parameters *c_p_* and *c_h_*. The error threshold 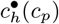 between weak and strong control (WC, SC) is marked by a yellow line.

In the SC regime 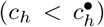, control maintains an evolved trait Γ* > 0 that is beneficial to host and pathogen. In the WC regime 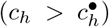, control is too weak to maintain the evolved trait and the pathogen reverts to the wild type (Γ_wt_ = 0). Fig. 3B shows the control efficiency

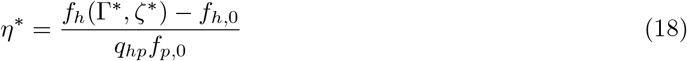

as a function of the cost parameters (*c_p_*, *c_h_*). In this case, *η*\* is a decreasing function of both parameters; i.e., host and pathogen cost impede evolutionary control. The error threshold 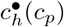 marks the transition between *η*\* > 0 in the SC regime and *η*\* = 0 in the WC regime. At the transition, *G*\* and *ζ*\* switch, while *f_p_* and *f_h_* remain continuous (Fig. S1).

### Control dynamics

How can the stationary level control value be reached by a dynamical control trajectory, and how can this trajectory be optimized? To address these questions, we have to specify the dynamics of the control amplitude. This can be a local update rule of the form

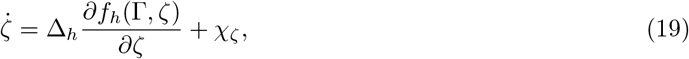

with white noise of mean 0 and variance ⟨*χ*_*ζ*_ (*t*)*χ*_*ζ*_ (*t*′)〉 = *D*_*ζ*_ *δ*(*t − t*′), similar to the stochastic trait dynamics given by Eqs. (1), (3). Local update can be realized in an evolutionary host system, where *ζ* is a quantitative trait with heritable variation Δ_*ζ*_ = 2*N*_*h*_*D*_*ζ*_ evolving in the fitness seascape *f_h_*(Γ(*t*), *ζ*)) [12]. Alternatively, *ζ* can be updated by a greedy protocol that finds the global maximum of the host fitness given the current pathogen state, *ζ*_max_(*t*) = arg max_*ζ*_ *f_h_*(Γ(*t*), *ζ*). In a complex landscape *f_h_*(Γ, *ζ*), global maximization by Darwinian evolution becomes mutation-limited and slow, unless the host population is large and the relevant range of *ζ* variants is contained in its standing variation. These dynamics can be implemented more rapidly by regulation: the pathogen load *L_p_* and the control amplitude *ζ* are positive and negative input variables, respectively, into a regulatory or metabolic network that produces an approximate *ζ*_max_ as output. A more sophisticated global procedure is to determine the optimal control protocol *ζ*\*(*t*) by maximizing the time integral of the host fitness, ∫ *f_h_*(Γ(*t*), *ζ*(*t*)) *dt*, over all control paths (Γ(*t*), *ζ*(*t*)) linking a given initial trait Γ_0_ with the equilibrium point, lim_*t*→∞_ Γ(*t*) = Γ*. In Methods, we show that this problem reduces to a local optimization condition for control protocols *ζ*(*t*) that can be solved in a straightforward way. Path optimization requires knowledge of the dynamical rules, as well as computation of the future dynamics by methods of control theory; this procedure can inform control designed by humans but hardly be realized by evolution or regulation.

These update rules generate a complex spectrum of control dynamics, the biological consequences of which will be discussed below. Over a broad parameter range of ecological control, the optimal stationary protocol (Γ*, *ζ*\*) given by Eq. (13) is a stable equilibrium of the local host-pathogen dynamics, which implies (Γ*, *ζ*\*) = arg max_*ζ*_ *f_h_*(Γ*, *ζ*) (Methods). As shown in Fig. 4A, a locally updated stochastic path (*D* > 0, *D*_*ζ*_ > 0; blue) with a sufficiently close starting point gets localized to (Γ*, *ζ*\*); the corresponding deterministic path (*D* = *D*_*ζ*_ = 0; orange) converges to (Γ*, *ζ*\*). The greedy-optimized (green) and the path-optimized (brown) protocol converge to the same point. All four protocols have only transient differences in their time-dependent efficiency (Fig. 4B). We conclude that the ecological control equilibrium can be dynamically reached and maintained in a robust way. In contrast, the optimal stationary protocol (Γ*, *ζ*\*) of our evolutionary control model is not a fixed point of the deterministic host dynamics. Hence, it cannot be reached or maintained by an evolutionary host system operating on local control update but requires path-optimized control (Fig. S2). Protocols based on global optimization can also be drawn towards an unstable or metastable fitness maximum (Methods, Fig. S2). Most strikingly, as the next example shows, optimal control can become time-dependent even if the control objective (maximizing the time-averaged host fitness) is not.

**Figure 4:**
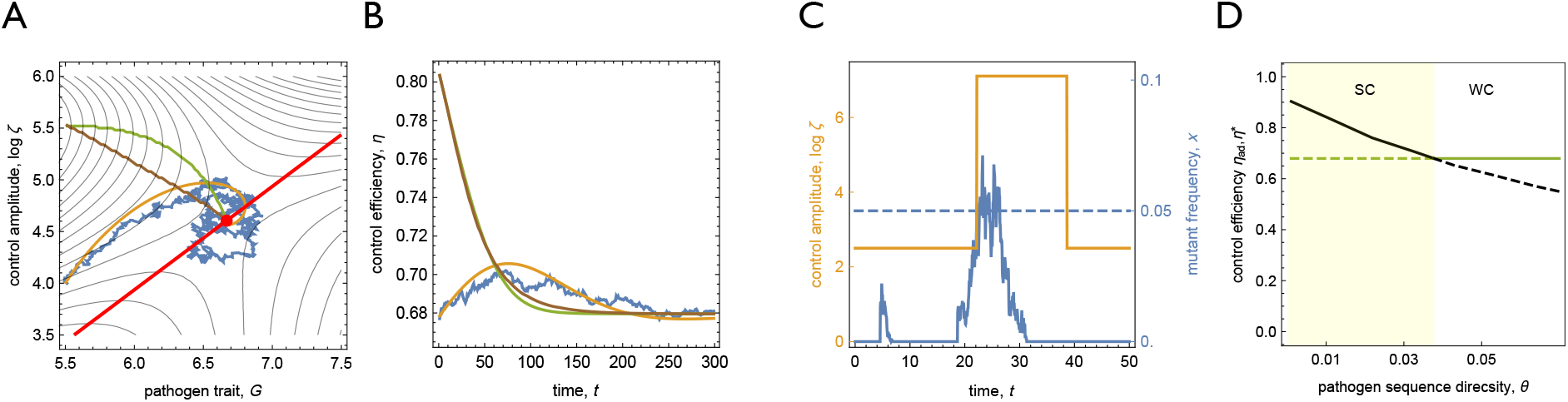
Control dynamics. (A) Approach to equilibrium for different protocols of ecological control: time-dependent control amplitude *ζ*(*t*) and resistance trait *G*(*t*) under local stochastic (blue), local deterministic (orange), greedy-optimized (green), path-optimized (brown) control dynamics. (B) Time-dependent control efficiency *η*(*t*) for the same protocols. (C) Adaptive ecological control: time-dependent protocol *ζ*(*t*) with baseline amplitude *ζ*_base_, boost amplitude *ζ*_res_, and boost duration τ_res_ (yellow); pathogen escape mutant frequency *x*(*t*) (blue), using a detection threshold *x*(*t*) > 0.05 to initiate rescue boost (dashed blue line). (D) Control efficiency of the optimal adaptive control protocol, 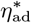 (black), as a function of the pathogen sequence diversity *θ*, together with the efficiency of the optimal stationary control, *η*\* (green), solid line shows preferred choice given *θ*. For *θ* < *θ*_*c*_ (yellow shading), adaptive control is more efficient than stationary control. Model parameters as in Fig. 2.

### Adaptive control

The phase diagram of Fig. 3A applies to sufficiently large pathogen populations, which reach an equilibrium with evolved resistance by frequent escape mutations. In smaller populations, these mutations become rare and the pathogen will stay in a metastable wild type over extended periods. Provided we can detect escape mutants at low frequency, we can keep the bulk pathogen population in the wild type by a two-state adaptive protocol of ecological control (Fig. 4C). As long as no escape mutant is detected, we apply a baseline control amplitude *ζ*_base_ to jointly curb pathogen load and escape rates. This task has to factor in mutational targets and selection coefficients over a spectrum of escape mutants with different trait values *G*. Once an escape mutant is detected above a threshold frequency *x*_0_, we apply a rescue boost with a larger amplitude *ζ*_res_ over a period τ_res_ to drive that mutant to loss. The efficiency of this protocol takes the simple form *η*_ad_ = *η*_base_ − *θ δ*_res_, which depends on the pathogen sequence diversity *θ* = 2*μN*_*e*_ (Methods). It is determined by the baseline efficiency in the absence of escape, *η*_base_, which often exceeds the equilibrium *η*\* given by Eq. (16), and the average efficiency cost per rescue, *δ*_res_. Optimized protocols maximize *η*_ad_ by jointly tuning *η*_base_ and *δ*_res_ (Methods). In Fig. 4D, we compare the efficiency of adaptive and equilibrium control. For *θ* < *θ*_*c*_ = (*η*_base_ − *η*\*)/*δ*_res_, i.e., at sufficiently low mutation rates or in sufficiently small populations, adaptive control outperforms stationary control. Alternative adaptive protocols, which include temporarily suspending control [23] or switching to a secondary defence against escape mutants, depend on *θ* in a similar way. Hence, all of these adaptive protocols strengthen control in small populations.

## Discussion

In this paper, we solve the optimization problem for a minimal model of pathogen control operating on realistic bio-molecular host-pathogen interactions. Specific features of host and pathogen biology enter the eco-evolutionary control theory of this system at several stages. In particular, the control mechanisms are based on the physiological effects of host-pathogen interactions – here, antibody-antigen binding –, and the objective functions of control, Eqs. (11) and (12), account for the fitness effects of these interactions. As we have shown, these biological features strongly impact the eco-evolutionary control dynamics and the efficiency of optimized protocols.

A salient feature of eco-evolutionary control is the emergence of high- and low-efficiency control regimes separated by an error threshold (Fig. 3). This behavior can be traced to the nonlinearities in the Hill function of antibody-antigen binding, Eq. (10). First, binding-mediated control has a diminishing-return leverage, which is bounded by the pathogen selection coefficient *q_ph_*. Second, the control cost depends exponentially on the chemical potential (log *ζ*), which determines the evolved pathogen state Γ_*e*_ (Fig. 2). Hence, the minimum-leverage control (8) has a moderate cost in the SC regime, but is too expensive or mechanistically impossible in the WC regime. Because nonlinearities of this form are a generic feature of bio-molecular interactions, we expect error thresholds to emerge also in more complex models of pathogen control.

In biomedical applications of pathogen control, it is crucial to predict the likely efficiency of control prior to any intervention. Our results show that few, independently measurable cost parameters can inform such estimates. The pathogen cost parameters (*c_p_* and *q_ph_* in the minimal model) are routinely measured in dosage-response assays [24]. The control cost parameter *c_h_* can, for example, be estimated as the metabolic production cost per unit of antibody [25, 26]. The arguably most case-specific parameter is the pathogen cost to the host. This parameter can be measured by fitness assays in microbial systems, but is more complex to assess for human hosts.

An equally important issue for control by humans and by natural hosts is optimizing the control dynamics in tune with the monitoring and computation capabilities of the host system. Local-update dynamics can efficiently be realized by Darwinian evolution (i.e., by variation of and selection on control molecule levels), whereas global-optimization control strategies depend to various degree on computation. In the minimal model, we find that local-update dynamics can reach the optimal stationary protocol (i.e., the point of maximal host fitness) for ecological control, but misses this point in the case of evolutionary control. Protocols based on regulation can, in principle, circumnavigate fitness valleys and implement an approximate maximization of the host fitness; however, signalling and regulatory networks themselves generate an additional cost to the host. Computation can further improve control, but its success is limited by the ability to predict pathogen evolution. A case in point is vaccine selection for human influenza based on predictive analysis [8]. It will also be interesting to analyze mechanisms and efficiency of control by “computation” in natural, non-human hosts.

Adaptive control, a particularly interesting strategy based on monitoring, realizes the time-independent objective to maintain a controlled pathogen state by a time-dependent protocol that responds to escape mutations [27]. In the control theory literature, this class of protocols is known as closed loop control [14]. Here we show that adaptive control can outperform the optimal stationary protocol (Γ*, *ζ*\*), which, for ecological control, is the Nash equilibrium of the host-pathogen fitness landscape in Eqs. (11), (12). Two conditions for efficient adaptive control emerge from our analysis: there is a metastable point of high host fitness (undercutting the minimum-leverage relation (8)), and escape mutants have a substantial intrinsic cost (generating positive selection for the wild type during rescue boosts). The optimal adaptive protocol explicitly depends on selection, mutation rate, and population size of the pathogen, highlighting the tight link of eco-evolutionary control theory to the underlying ecological dynamics and population genetics. This implies that eco-evolutionary control theory needs to factor in both absolute and relative fitness [28].

In summary, this paper establishes a conceptual framework for eco-evolutionary control and infers robust features of bio-molecular control, based on a minimal model of host-pathogen interactions with stationary control objectives. Our underlying leverage calculus extends to non-equilibrium control problems, but temporal constraints or explicitly co-evolutionary dynamics [12] generate different objective functions. Several complexities of biomedical control problems are not captured by the minimal model. For example, the human immune system has a complex antibody repertoire, and pathogens often have multiple antigenic binding sites (microbial and viral epitopes, cancer neoantigens). This generates multiple antibody-antigen interactions, i.e., multiple potential control channels with independent control amplitudes, leverage, and cost parameters. Conversely, pathogens with a high mutation rate often have multiple channels of escape mutations from a given antibody [29]. Consequently, the fitness landscapes of host and pathogen live in a multidimensional parameter space; an appropriate control dynamics is to navigate this space towards high-efficiency protocols. An important example is the dynamics of broadly neutralizing and specific antibodies for HIV [11, 13]. Another layer of complexity arises for pathogens embedded in microbiota, which are multi-species systems with a tightly connected ecology. The control of a given pathogen can perturb the entire microbiota, generating cross-resistance of multiple pathogens and complex collateral effects for the host [30]. In immune systems, the control of multiple pathogens requires a complex pattern of resource allocation [31]. Gearing up eco-evolutionary control theory to these systems is an important avenue for future work.

## Acknowledgments

We thank Armita Nourmohammad and Johannes Cairns for discussions. This work has been supported by Deutsche Forschungsgemeinschaft grants SFB 680 and SFB 1310 (to ML). We acknowledge the CSC–IT Center for Science, Finland, for computational resources.

## Methods

### 1. Eco-evolutionary control theory

Here we describe the probabilistic calculus for the evolution of a quantitative trait and the concurrent population dynamics in a given ecological niche and we derive minimum-leverage relations used in the main text.

#### Stochastic eco-evolutionary dynamics

These dynamical processes of a quantitative trait *G* and of the population size *N* are generically coupled by selection: fitness effects generated by the evolution of the trait change the population’s carrying capacity, which, in turn, affects the natural variation of the trait and the speed of evolution. The diffusion equations (1) – (4) used in the main text depend on the diffusion coefficients for the population mean trait Γ and the population size *N*, which enter Eqs. (3) and (4), and on the response coefficients in Eqs. (1) and (2) [1, 2]. The trait diffusion coefficient *D* is given by the increase of the heritable trait variance per generation by new mutations,

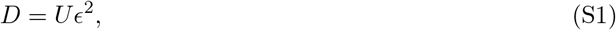

which depends on the total mutation rate at trait-encoding genomic sites, *U*, and the mean square trait effect of these mutations, *ϵ*^2^. The trait response to the entropic force (*dS*(Γ)/*d*Γ) equals *D*, which ensures the neutral trait dynamics leads to the correct equilibrium trait distribution given by Eq. (S4) below. The trait response to selection depends on the response coefficient

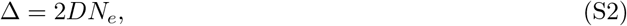

which equals the equilibrium expectation value of the heritable trait variance. Over a broad range of evolutionary conditions, Eq. (S2) can be regarded as a defining relation for the effective population size *N_e_* [2], similar to the relation *π* = 2*μN_e_* that links the sequence variance at neutral sites to the point mutation rate *μ* [3]. The effective population size *N_e_* given by Eq. (S2) is also broadly related to the coalescence time, *τ*_*c*_ = 2*N_e_* [4, 5]. The relation of *N_e_* to the census population size *N* depends on the system and its evolutionary mode; some relevant cases are listed below.

The population size dynamics given by Eqs. (2), (4) with the diffusion constant *N/*2 can be derived from a simple birth-death model. We consider a Poisson process with birth rate *b*(Γ,*N*) and death rate *d*(Γ,*N*), in which the reproductive rate

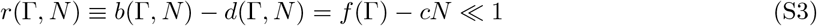

depends on the population size and the niche constraint parameter *c*, and the constraint *b*(Γ,*N*) + *d*(Γ, *N*) = 1 is satisfied. To leading order in *r*, this model generates the processes 1 → 0 with probability *p* = 1/4, 1 → 1 with *p* = 1/2, 1 → 2 with *p* = 1/4, leading to a diffusion constant as given in Eq. (4). The expectation value of the population size, i.e., the carrying capacity, is 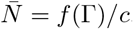. Other models of this class differ in their detailed birth and death rates but generate a similar diffusive dynamics of large population sizes (i.e., in a Taylor expansion about 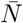) [6, 7].

#### Neutral eco-evolutionary equilibrium

In the special case of neutral evolution, i.e., for a trait- and time-independent reproductive rate *f* (Γ, *t*) = *f*_0_, the evolutionary dynamics given by Eqs. (1) – (3) has an equilibrium distribution

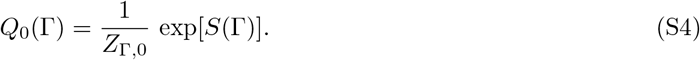

According to this distribution, the probability of a given Γ value is proportional to the density of sequence states mapped onto that value. Here we assume equiprobability of sequence states under neutral evolution; i.e., we ignore sequence composition biases at trait loci, which are usually small and irrelevant for this paper.

Similarly, the population dynamics given by Eqs. (2) – (4) has an equilibrium distribution

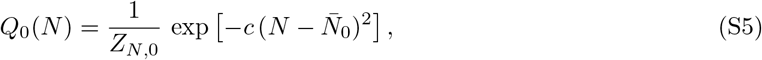

which describes stable fluctuations of the population size around the carrying capacity of the ecological niche,

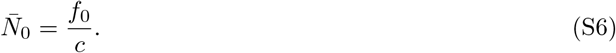

Here we assume that this carrying capacity is large enough so that the probability of fluctuations leading to extinction of the population can be neglected [8].

#### Eco-evolutionary stationary states under selection

The evolutionary dynamics of Eqs. (1) – (3) can be integrated in any time-independent fitness landscape *f* (Γ). The equilibrium distribution

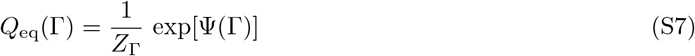

can be written in terms of the *free fitness*

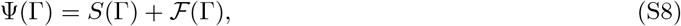

which is the sum of the entropy *S*(Γ) defined in Eq. (S4) and the fitness potential

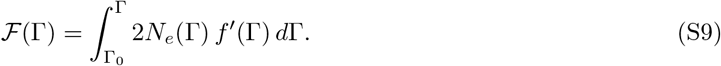

The integral on the r.h.s. contains an arbitrary reference point Γ_0_, variations of which can be absorbed into the normalization factor *Z*_Γ_. The fitness potential (S9) takes into account that the effective population size generically depends on the trait value, *N_e_* = *N_e_*(Γ), which generates an inhomogeneous response to selection. The form of this response depends on the evolutionary mode of the system. (i) If the coalescence process is dominated by genetic drift, the coalescence time exceeds the relaxation time of the population size in a stable population, *τ*_*c*_ ≫ *τ*_*N*_, as shown in the next paragraph. In this case, we can often approximate 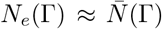; hence, the evolutionary dynamic decouples from the instantaneous census population size and becomes autonomous. If the evolutionary range of Γ generates only small relative differences in absolute fitness, 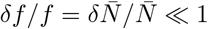, we can further approximate *N_e_* by a constant. The fitness potential then reduces to the standard form of a reduced fitness,

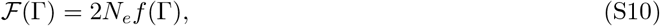

(ii) Under clonal evolution in large asexual populations, the coalescence process is dominated by interference effects. This generates effective population sizes with a very weak dependence on *N* ; for example, *N_e_* ~ (log *N*)^1/3^ in the travelling fitness wave model [5]. The dominant contribution to the integral in equation (S9) then comes from variations in *f* (Γ); hence, for the leading-order estimates used below, we can still use the approximation (S10). (iii) For extreme non-equilibrium processes, which involve likely trajectories with population sizes far from the carrying capacity, we expect the evolutionary process to couple to the path-dependent census population size. The fluctuation statistics of this regime is more involved and falls outside the scope of this paper.

The population dynamics of Eqs. (2) – (4) reaches a simple conditional equilibrium distribution

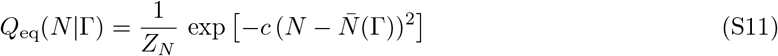

with the trait-dependent carrying capacity

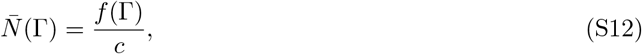

if the relaxation time of the population size, *τ*_*N*_, is much smaller than the coalescence time, *τ*_*c*_ = 2*N_e_*. This condition is self-consistently fulfilled for stable populations with coalescence dominated by genetic drift. To show this, we estimate the coalescence time in terms of the time-averaged census population size at a given trait value, 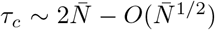. The relaxation time 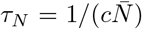 follows directly from Eq. (2). Together, the condition for time separation between population dynamics and evolution reads

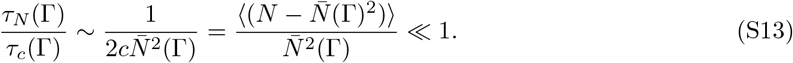

This relation involves the variance of the population size scaled by the squared carrying capacity, which is small for stable populations far from stochastic extinctions.

The distributions (S7) and (S11) can be combined into an equilibrium distribution of the joint eco-evolutionary dynamics,

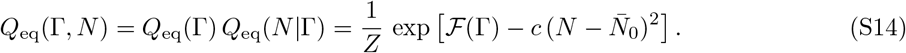

On the other hand, clonal evolution under strong interference selection generates shorter coalescence times *τ*_*c*_, which can become comparable to *τ*_*s*_. In that case, the stationary distribution *Q*(Γ,*N*) no longer has the simple equilibrium form (S14); it becomes a non-equilibrium state with a nonvanishing probability current. However, the marginal trait distribution retains the equilibrium form *Q*_eq_(Γ) given by Eq. (S7), as long as the trait dynamics remains (approximately) autonomous.

#### Non-equilibrium fluctuation theory and evolutionary flux

The stochastic dynamics given by Eqs. (1) – (4) can be applied to estimate the probability of adaptive eco-evolutionary processes in a generically time-dependent fitness seascape *f* (Γ, *t*). A realization of this process in a given population maps an evolutionary path (**Γ**, **N**), which is defined by continuous functions (Γ(*t*),*N* (*t*)) connecting an initial population state (Γ_*i*_, *N_i_*, *t_i_*) to a final state (Γ_*f*_, *N_f_*, *t_f_*) (*t_i_* ≤ *t* ≤ *t_f_*). The relevant non-equilibrium fluctuation theory is developed in refs. [9, 10, 11, 12] and reviewed in ref. [13]; a detailed discussion of evolutionary fluctuation calculus is given in ref. [14]. Here we first evaluate the probability (density) 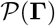 of a marginal trait path **Γ** evolving in a fitness seascape *f* (Γ, *t*), which is the product of the probability of the initial state at time *t_i_* and the conditional probability of the path for given initial state,

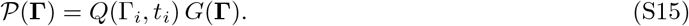

This forward path probability is to be compared with the probability 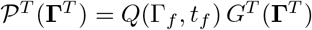 of the time-reversed (backward) path Γ^*T*^, which is defined by the map Γ^*T*^ (*t*) ≡ Γ(*t_f_* + *t_i_* − *t*) evolving in the time-reversed fitness seascape *f^T^* (Γ, *t*) ≡ *f* (Γ, *t_f_* + *t_i_* − *t*) (*t_i_* ≤ *t* ≤ *t_f_*). A key result of non-equilibrium fluctuation theory expresses the ratio of the forward and the backward conditional probabilities in the form

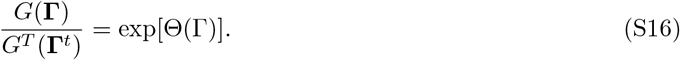

Here Θ(**Γ**) is the evolutionary flux of the forward path,

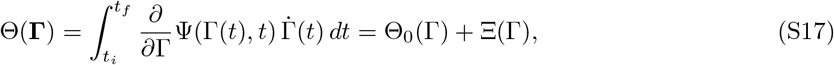

where

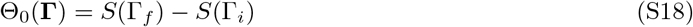

is the neutral flux and

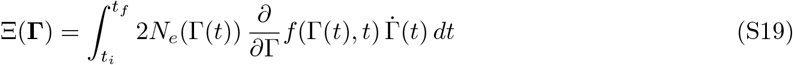

is the scaled fitness flux of the forward path **Γ**. The relations (S16) – (S19) can be derived by a straight-forward generalization of the analogous result in ref. [14]. They are valid as long as the evolutionary dynamics can be written in an (approximately) autonomous form, i.e., with an effective population size *N_e_*(Γ). As discussed above, this includes evolutionary parameter regimes with coalescence dominated by genetic drift or by interference selection.

In the special case of a time-independent fitness landscape, the evolutionary flux Θ(**Γ**) reduces to the difference in free fitness between the final and initial state of the path **Γ**,

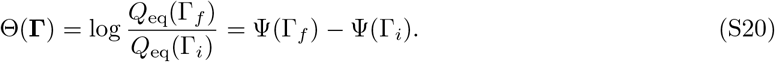

This result follows by comparison of Eqs. (S18), (S19) with Eq. (S7) and expresses detailed balance of the evolutionary equilibrium state; i.e.,

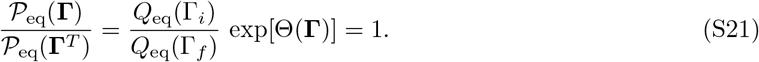

Furthermore, in the regime of fast population dynamics (*τ*_*N*_ ≪ *τ*_*c*_), Eq. (S16) can be generalized to full eco-evolutionary trajectories in a straightforward way,

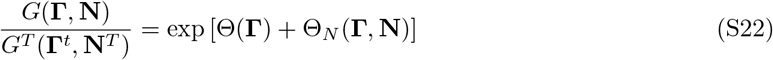

with the population size flux

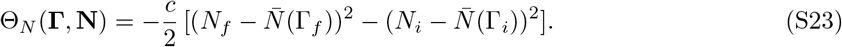

#### Fitness flux theorem

An immediate consequence of the fluctuation relation (S16) is the identity

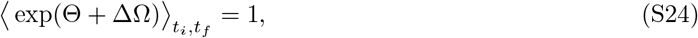

where Θ(**Γ**) is the evolutionary flux of a given path **Γ**defined by Eqs. (S16) – (S19),

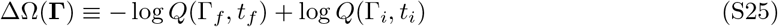

is the difference in local entropy between the final and the initial state of the path **Γ**, and 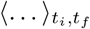 denotes averages over the ensemble of paths **Γ** in the time interval (*t_i_*, *t_f_*). Following standard fluctuation theory, this identity is proved by using Eqs. (S15) and (S16) to rewrite the path weight, 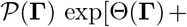 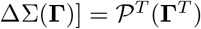, and integrating over the path ensemble; Eq. (S24) then reduces to the normalization condition 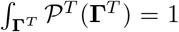. By defining the difference in local relative distance to the neutral ensemble between the final and the initial state of the path **Γ**,

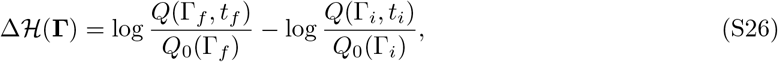

we can rewrite Eq. (S24) in an equivalent form containing the scaled fitness flux,

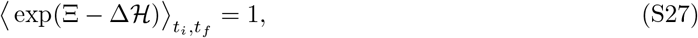

Details of definitions and derivations are given in ref. [14]. The flux identities (S24) and (S27) are valid for any evolutionary process given by stochastic equations of the form (1) – (4). Importantly, they do not depend on specific properties of the initial and the final state; that is, they are valid over the full duration (*t_i_*, *t_f_*) of the process or over any subperiod 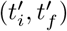.

By applying Jensen’s inequality to Eq. (S24), we obtain the inequality

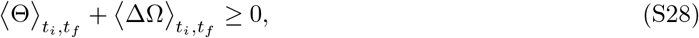

which is again valid for any (sub-)interval 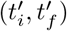 of the process. Here and in the following relations, equality applies to equilibrium and strict inequality to spontaneous processes (i.e., non-equilibrium processes that take place with a finite speed). In the special case of a time-independent fitness landscape, this relation can be rewritten in terms of free fitness differences,

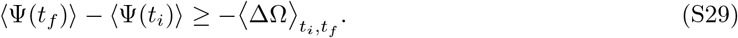

In the following, we will use a deterministic form of the inequality (S28),

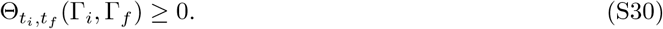

This form emerges in two ways: (i) For traits with a sufficiently large number of encoding genome sites, the distributions Ω(Γ, *t_i_*) and Ω(Γ, *t_f_*) become peaked around the maximum-likelihood values 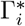 and 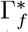, respectively. Integrating over these distributions, neglecting the subleading contribution from the ΔΩ term, and dropping the star notation leads to the form (S30), in analogy with the thermodynamic limit. In practice, the form (S30) provides already a reasonable approximation for transcriptional protein-protein binding traits that are encoded in ~ 10 - 20 sequence sites. (ii) A similar dominance of the Θ term in equation (S28) arises in the regime of large *N_e_*, which corresponds to large scaled selection amplitudes 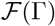. The relation (S30) says the trait moves on average uphill on the instantaneous free fitness landscape Ψ(Γ, *t*), which is the sum of the time-dependent fitness potential 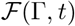 and the Boltzmann entropy *S*(Γ), defined as the log number of sequence states with trait value Γ. For time-independent fitness landscapes, it reduces to

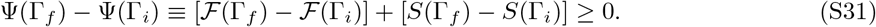

#### Minimum leverage relations

The evolutionary flux calculus developed here can be used to map the efficiency of a wide range of eco-evolutionary control processes. If the uncontrolled system is governed by a fitness landscape *f_n_*(Γ) and the controlled system by a fitness seaor landscape *f* (Γ, *t*) = *f_n_*(Γ)+ *f_c_*(Γ, *t*), selection and entropy generate the free fitness functions Ψ_*n*_(Γ) and Ψ(Γ, *t*), respectively, by Eqs. (S8) and (S9). Maintaining a controlled trait value Γ_*c*_ can be achieved by a stationary control landscape *f_c_*(Γ). A minimum-leverage condition for maintenance of local equilibrium Γ_*c*_ against another local equilibrium Γ_0_ is given by Eq. (7) of the main text, Ψ(Γ_*c*_) − Ψ(Γ_0_) ≥ 0. This sufficient condition follows directly from the relation (S31); it is also necessary if escape mutations would otherwise generate a spontaneous evolutionary path from Γ_*c*_ to Γ_0_ on the time scale of the control process. If escape mutations occur at a sufficiently low rate, metastable values Γ_*c*_ can be maintained by a lower minimum leverage of control. In the main text, we use the relation (8) to map phase diagrams delineating strong and weak control in large populations (where escape mutations are assumed to be frequent), and we discuss an example of adaptive control maintaining a metastable equilibrium in small populations (with rare escape mutations).

### 2. Application to pathogen control

Here we derive the analytical solution of stationary control, and we detail the different control dynamics used in the main text and in Fig. 4. These include local deterministic and stochastic, greedy-optimized, path-optimized, and adaptive protocols, which are evaluated in the fitness landscapes of Eqs. (11), (12).

#### Pathogen equilibrium states

In the parameter regime of interest, the pathogen fitness landscape (11) has two local fitness maxima: (i) The wild type Γ = 0 (vertical line in Fig. 2) is assumed to be a boundary; i.e., the trait evolution generates only values Γ ≥ 0. (ii) The evolved state Γ_*e*_(*ζ*) (inclined line in Fig. 2) exists in the regime

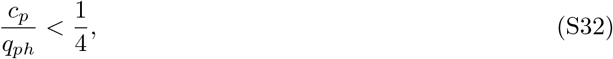

which provides a bound to the pathogen cost. Maximization of *f_p_*(*G*, *ζ*) with respect to *G* determines two characteristics of the evolved state. (i) The trait value Γ_*e*_ is shifted by an amount

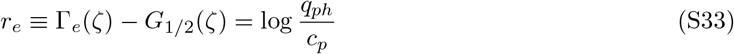

with respect to the half-binding point *G*_1/2_(*ζ*) (which equals the trait value that the IC50 value *ζ*). (ii) The error *q_e_*, which is defined as *q_e_* = *P*_bind_ for ecological control (*ϵ* = +1) and as *q_e_* = 1 − *P*_bind_ for evolutionary control (*ϵ* = −1), is given by

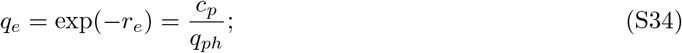

here we have used the exponential asymptotic form of the binding probability (10), which is a good approximation for parameters satisfying Eq. (S32). These characteristics determine trait and fitness of the evolved state,

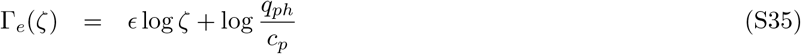

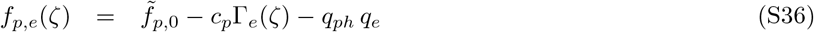

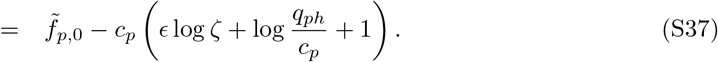

with 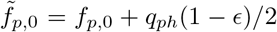. The fitness ranking, i.e., the evolutionary stability, of the evolved state and the wild type depends on the control amplitude *ζ* (stable/metastable pathogen states are shown as solid/dashed line segments in Fig. 2). The rank switch, which marks the loss of the evolved trait, occurs at a value *ζ*_*l*_ determined by the condition *f_p,e_*(*ζ*_*l*_) = *f_p_*(*ζ*_*l*_, 0) (horizontal line in Fig. 2). This determines the crossover point

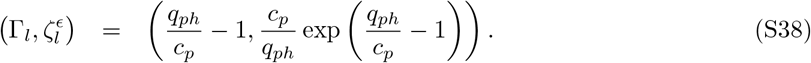

#### Stationary ecological control

We compute the point of optimal stationary control, (Γ*, *ζ*\*), by evaluating the host fitness with the pathogen at its conditional stable equilibrium, *f_h_*(Γ*(*ζ*), *ζ*), and maximizing this function with respect to *ζ*. For ecological control in the strong-coupling (SC) regime, the pathogen is in the wild type Γ_wt_ = 0; in the weak-coupling (WC) regime, the pathogen is in the evolved state Γ_*e*_(*ζ*) given by Eq. (S37). We obtain the equilibrium point (Γ*, *ζ*\*) as given by Eq. (14); this point has the pathogen fitness

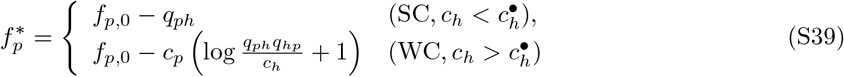

and the host fitness

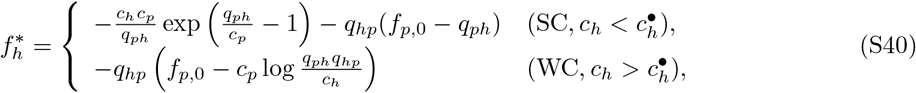

where we have used the approximation *P*_bind_ ≈ 1 in the wild type. The efficiency of control is obtained by inserting 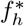 into the definition (S41),

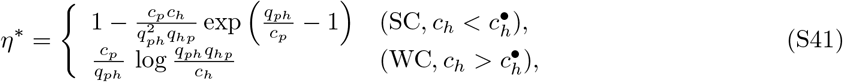

In Fig. S1AB, the optimal control coordinates (Γ*, *ζ*\*), the resulting fitness values 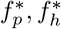, and the control efficiency *η*\* are plotted as functions of the control cost *c_h_*. The transition between the SC and WC control regimes is determined by the condition Γ_*e*_ = Γ _*l*_, from which we obtain the transition line 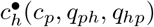 given by Eq. (15).

#### Stationary evolutionary control

We again maximize the host fitness *f_h_*(Γ*(*ζ*), *ζ*) with respect to *ζ* with an evolved pathogen trait Γ_*e*_(*ζ*) in the SC regime and a wild type trait Γ_wt_ = 0 in the WC regime. We obtain the optimum-control point

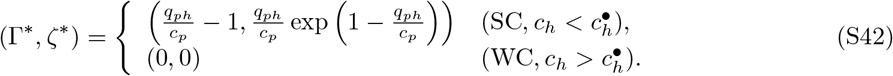

The optimal SC control is now at (Γ* = Γ_*l*_, *ζ*\* = *ζ*_*l*_) given by Eq. (S38), where the pathogen has the maximum stable Γ_*e*_ value. The stationary point has the pathogen load

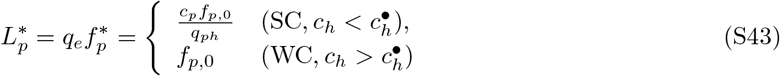

with *q_e_* = (1 − *P*_bind_) given by Eq. (S34), and the host fitness

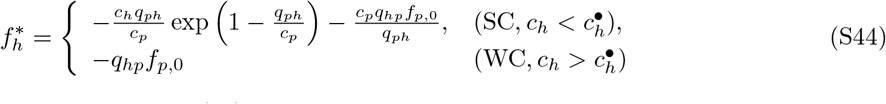

This determines the efficiency of control (19),

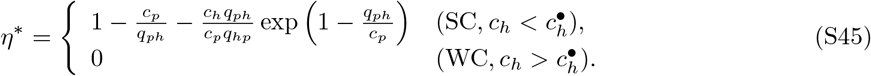

In Fig. S1CD, the optimal control coordinates (Γ*, *ζ*\*), the resulting fitness values 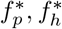, and the control efficiency *η*\* are plotted as functions of the control cost *c_h_*. The transition between the SC and WC control regimes is determined by the condition 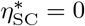, from which we obtain the transition line 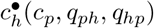 given by Eq. (18).

#### Local control dynamics

The stationary point (*G*\*, λ*) given by Eq. (13) is an optimal time-independent control protocol: it maximises the host fitness, given that the pathogen is at the point *G*\*(*ζ*) of its conditional fitness maximum for any given control amplitude *ζ*. Here we analyze the evolutionary stability of this protocol under local dynamics of pathogen and host, as given by Eqs. (1) and (20). We use the deterministic form

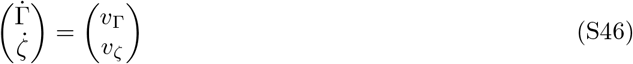

with the velocity field

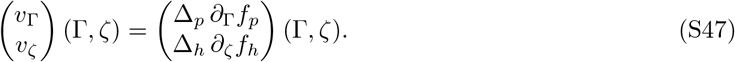

Both components of the deterministic dynamics (S46) take the form of Breeder’s equation. Here we have omitted the entropic force contained in Eq. (1), which is appropriate for large pathogen populations. (i) In our model of ecological control, (*G*\*, λ*) is a fixed point of the deterministic dynamics,

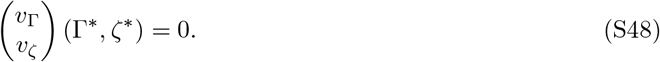

**Figure S1:**
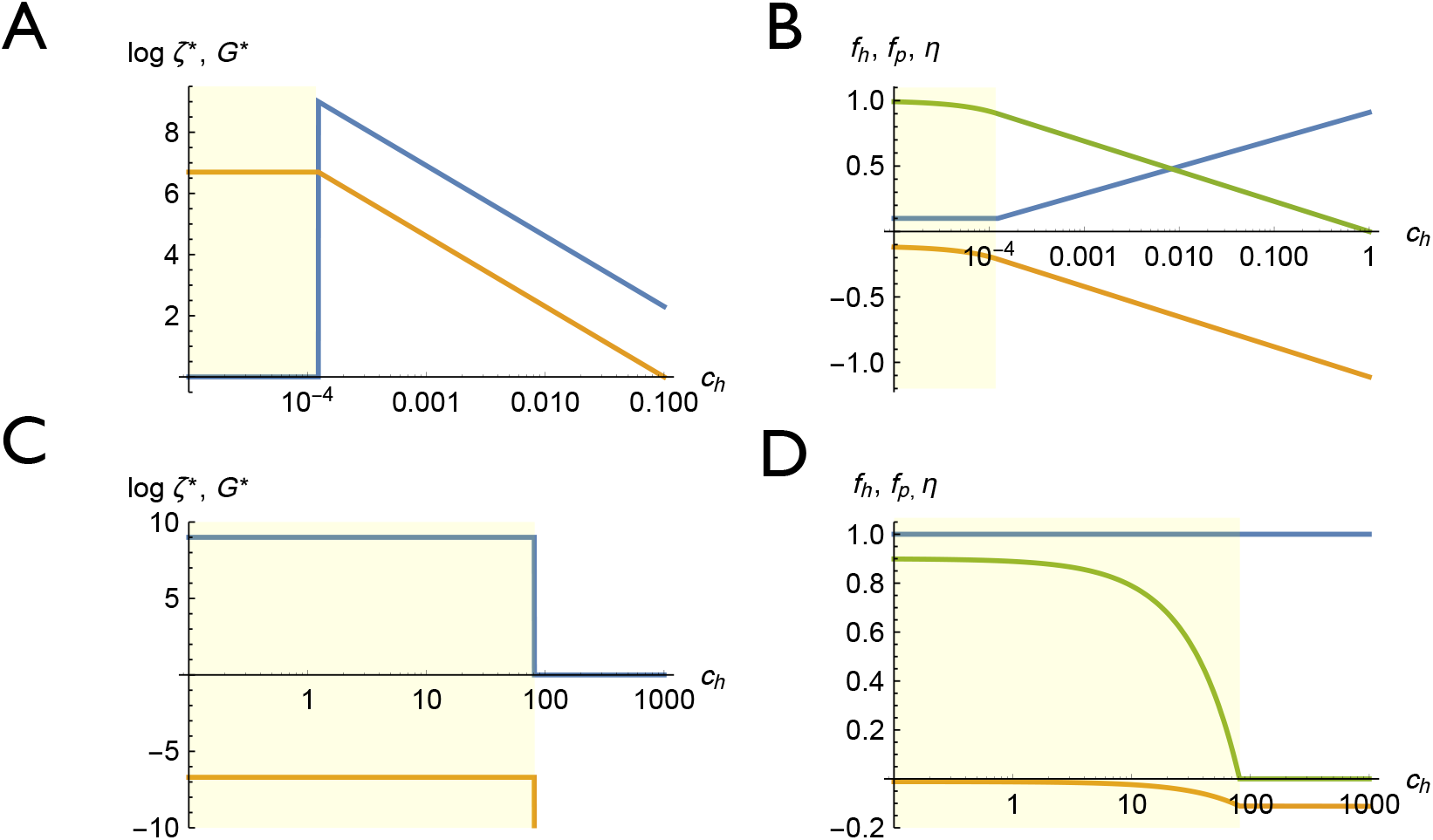
Stationary control protocols. (A, B) Ecological control, (C, D) evolutionary control. (A, C) Equilibrium log control amplitude, log *ζ*\* (orange), and pathogen trait, *G*\* (blue), as functions of the control cost, *c_h_*. Yellow shading marks the strong control regime. (B, D) Host fitness, 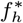 (orange), pathogen fitness, 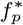 (blue), and control efficiency, *η*\* (green), as functions of *c_h_*. Model parameters: *f_p_* = 1, *q_ph_* = 1/*q_ph_* = 0.9, *c_p_* = 0.09 (ecological control); *f_p_* = 1, *q_ph_* = 1/*q_ph_* = 9, *c_p_* = 0.9 (evolutionary control).

Using Eqs. (11) and (12), the stability of this fixed point can be expressed in terms of the binding probability,

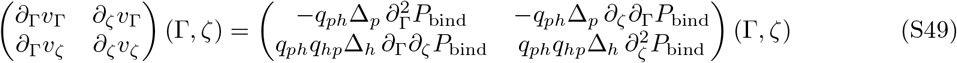

with *p*_bind_(Γ, *ζ*) given by Eq. (10). It is straightforward that this matrix is negative-definite in the parameter regime of interest; see Eq. (S32). Hence, the optimal stationary-control point (*G*\*, λ*) is a stable fixed point of the deterministic evolutionary dynamics given by Eq. (S47). As shown in Fig. 4AB, a deterministic control path (Γ(*t*), *ζ*(*t*)) starting in the vicinity (orange line) converges to (*G*\*, λ*); a corresponding stochastic path (blue line) gets localized to the same point. (ii) In our model of evolutionary control, (*G*\*, λ*) is not a fixed point of the deterministic host-pathogen dynamics. In this case, the velocity field (S47) has a component *v*_*ζ*_ (*G*\*, λ*) ≠ 0, i.e., the host can gain a transient fitness advantage by moving away from the optimal stationary amplitude *ζ*\*. Hence, the optimal stationary-control point (*G*\*, λ*) cannot be reached or maintained by a local control dynamics. As shown in Fig. S2B, deterministic and stochastic control paths (Γ(*t*), *ζ*(*t*)) starting in the vicinity of (*G*\*, λ*) move away from this optimal point.

#### Control dynamics with global optimization

In this paper, we consider two classes of control dynamics with global optimization: (i) Greedy protocols maximize the host fitness given the current pathogen state,

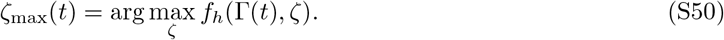

(ii) Path-optimized protocols maximize the time integral of the host fitness,

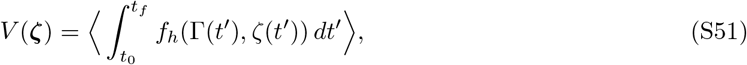

over a family of control paths *ζ* defined in the time interval (*t*_0_, *t_f_*). The stochastic process given by Eq. (1) generates a probability distribution over evolutionary paths **Γ**, and angular brackets denote expectation values in this path ensemble. We now discuss how path-optimized protocols can be computed for the host-pathogen control problems discussed in the main text. With given boundary condition Γ(*t*_0_) = Γ_0_ the optimization problem can be solved by a transfer matrix approach. We consider the fitness integral (S51) for the set of paths defined in a partial time segment (*t*, *t_f_*) (*t*_0_ < *t* < *t_f_*) and constrained to a pathogen initial state Γ(*t*) = *G*,

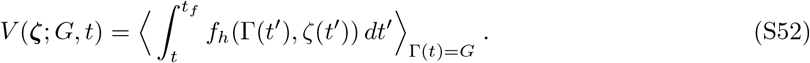

We define the conditional path-optimized protocol *ζ*\*(*G, t*) and the corresponding fitness integral *J* (*G, t*),

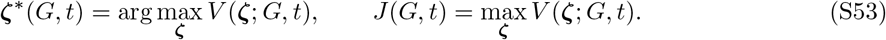

Given a diffusive pathogen dynamics of the form (1), (3), this conditional optimum can be shown to satisfy the local relation [15]

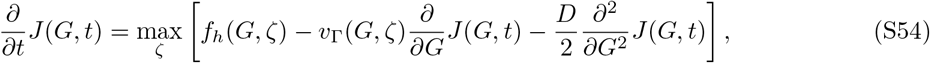

where *v*_Γ_(*G*, *ζ*) = Δ ∂_*G*_*f*_*p*_(*G*, *ζ*) is the deterministic pathogen velocity given by Eq. (S47) and *D* is the diffusion constant in Eq. (3). This relation is known as the Hamilton-Jacobi-Bellman equation for continuous-time stochastic control [15]. It can be used to compute the optimal solution *J* (*G, t*) and the associated optimal control path *ζ*\*(*G, t*) by recursion backwards in time. In the control theory literature, the path optimization problem is often formulated in terms minimization of the cost function (−*V* (*ζ*)), and the endpoint-conditioned optimum (−*J* (*G, t*)) is referred to as the cost-to-go function.

Here we are interested in the deterministic control problem (*D* = 0) with boundary conditions Γ(*t*_0_) = Γ_0_ and lim_*t*→∞_ Γ(*t*) = Γ*, i.e., for protocols that link an initial pathogen state Γ_0_ with the optimal stationary-control point (Γ*, *ζ*\*) given by Eq. (13) (this choice will be justified below). For large *t_f_*, solving Eq. (S54) leads to a stationary control profile where the optimal control *ζ*\*(*G*) does not depend explicitly on time and the l.h.s. of the equation takes a constant value 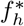,

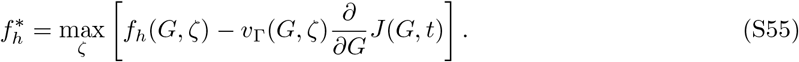

We maximize the right hand side with respect to *ζ* and solve for ∂_*G*_*J*,

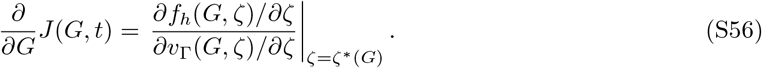

Using this relation, Eq. (S55) takes the simple form

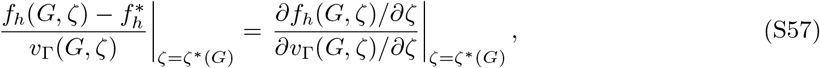

which is equivalent to the local path condition

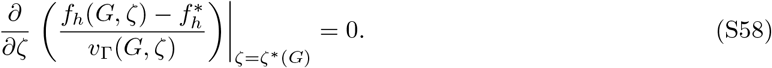

**Figure S2:**
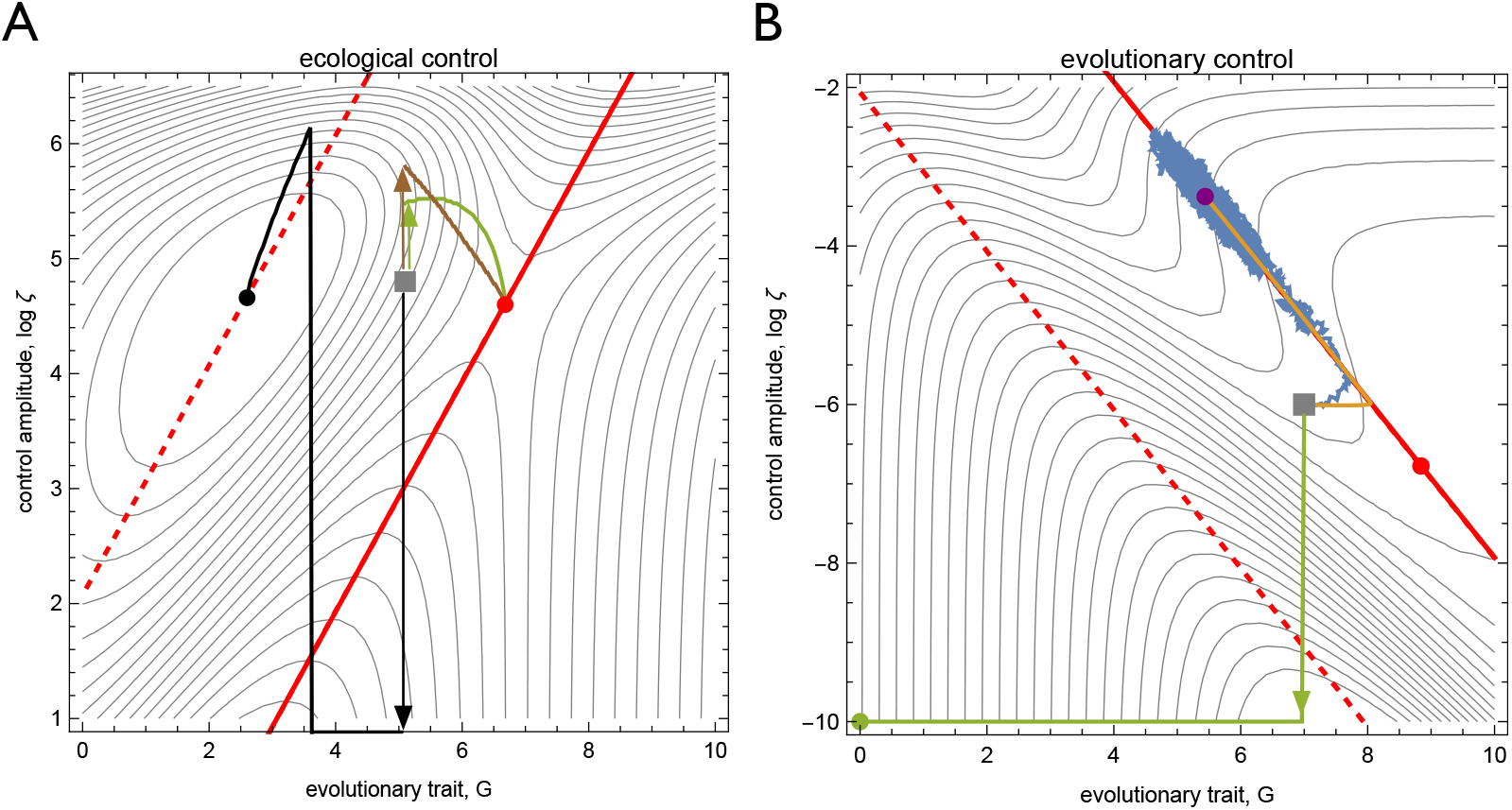
Control dynamics depends on update rule and boundary conditions. The figure shows different control paths starting from a common initial condition (grey square). (A) Ecological control. The globally path-optimized deterministic control path (black line, zero control shown as horizontal line) converges to an unstable fixed point (black dot) with disruptive selection on the pathogen trait. The path-optimized control path converging to the optimal stationary protocol (Γ*, *ζ*\*) (brown line, as in Fig. 4A) is obtained by maximization of the host fitness integral with an appropriate end point condition (see text). The greedy-optimized path with the same initial condition (green line) also converges to (Γ*, *ζ*\*). Host fitness is marked by contours; cf. Fig. 2B. (B) Evolutionary control. The point of optimal stationary control, (Γ*, *ζ*\*) (Γed dot), cannot be reached or maintained by local control dynamics. Deterministic and stochastic control paths with local update (orange and blue lines) move away from (Γ*, *ζ*\*) towards a stable fixed point (purple dot) with lower host fitness – i.e., a smaller control efficiency – than the SC optimal stationary protocol. The greedy-optimized control path (green line) sets *ζ* = 0 and drives the system to the WC fixed point (Γ_wt_ = 0, *ζ* = 0) (green dot), which also has lower host fitness than the SC optimal protocol (Γ*, *ζ*\*).

Substituting *G* = Γ(*t*), this relation describes the time-dependent control amplitude *ζ*\*(*t*) = *ζ*\*(Γ(*t*)) for path-optimized deterministic protocols converging to the optimal stationary point (Γ*, *ζ*\*) given by Eq. (13). Reducing the Hamilton-Jacobi-Bellman equation (S54) to a simple local optimization condition is possible because the host fitness function *f_h_* is not explicitly time-dependent.

In Fig. 4AB, we plot the greedy-optimized and path-optimized control paths (green and brown line) in the case of ecological control. Here, the path-optimized solution has been obtained by backward iteration of the Hamilton-Jacobi-Bellman equation (S54) within the trait range shown in Fig. S2A, or, equivalently, by evaluation of the local condition (S58). In Fig. S2A, we compare these paths to path-optimized control with a different boundary condition, 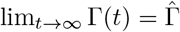, where

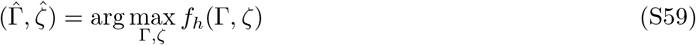

is the point of the absolute host fitness maximum in Fig. 2A. The deterministic control path converging to 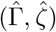 (black line in Fig. S2A) is a well-defined solution of the Hamilton-Jacobi-Bellman equation with an asymptotic host fitness 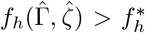; however, this solution has to be discarded on biological grounds. The fixed point 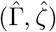 corresponds to a minimum of the pathogen fitness with respect to trait variation, 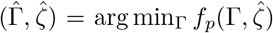 (the locus of these minima is shown as a dotted line in Fig. 2 and Fig. S2A). Hence, the pathogen population is under disruptive selection, i.e., mutants with 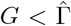 and with 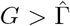 are under positive selection. These mutants cannot be simultaneously contained by control selection of the form *f_p_* ~ *P*_bind_ considered in this paper. We conclude that stationary control has to be in an evolutionarily stable pathogen state, as described by the condition (Γ*, *ζ*\*) = arg max_Γ_ *f_p_*(Γ, *ζ*\*) in Eq. (13). Path-optimized control protocols converging to such states are obtained by maximizing the host fitness integral, Eq. (S51), with appropriate boundary conditions. Fig. S2B shows the greedy-optimized control path (green line) in the SC regime of evolutionary control. This protocol sets *ζ* = 0, leading to a transient host fitness increase. The subsequent pathogen evolution drives the system to the WC fixed point (Γ_wt_ = 0, *ζ* = 0) (green dot), which has lower fitness than the SC optimal protocol (Γ*, *ζ*\*) given by Eq. (S42).

#### Adaptive control in finite populations

Adaptive protocols of pathogen control have a time-dependent control amplitude that depends on the presence of escape mutations in the pathogen population. Here we show that the time-average of the host fitness under adaptive control depends on the total mutation rate at trait-encoding sites, *u*_0_, and on effective population size *N_e_* of the pathogen population; in particular, adaptive control performs better than equilibrium control for small values of the diversity parameter *θ* = 2*N_e_u*_0_. We consider a specific class of adaptive protocols of ecological control that maintain a wildtype pathogen (Γ = 0) by a baseline control amplitude *ζ*_base_ and suppress escape mutants of effect *G* by rescue boosts of amplitude *ζ*_res_(*G*) and duration *τ*_res_(*G*). We label these protocols by their parameters (*ζ*_base_, *ζ*_res_, *τ*_res_), with boldface letters indicating functions of the trait *G*.

Assuming that escape mutants are detected at a low frequency *x*_esc_ and subsequently suppressed by the rescue protocol, they have a negligible effect on the mean pathogen trait, Γ < (1 − *x*_esc_)*G*_−_ + *x*_esc_*G* = *G*_−_ + *O*(*x*_esc_). This leads to simple expressions for the time-average fitness of pathogen and host,

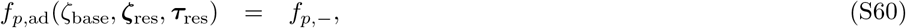

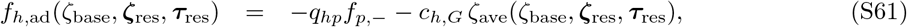

where *f*_*p*,−_ is the wildtype pathogen fitness given by equation (11) and *ζ*_ave_ is the time-averaged control amplitude. This amplitude depends on the control amplitudes and the pathogen dynamics in baseline and rescue mode. Under baseline control, we consider a pathogen population peaked at the wildtype trait value (*G*_−_ = 0), which a spectrum of trait mutations with rates *u*(*G*) = *μ ρ*(*G*), where *μ* is the mutation rate and *ρ*(*G*) is the mutational target for an effect of size *G*. A given mutant can trigger escape if it has a positive selection coefficient against the wildtype,

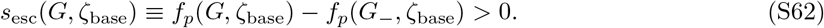

Hence, its establishment rate, which equals its fixation rate in the absence of a rescue control boost, is given by

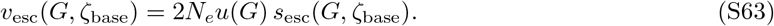

Suppressing this mutation requires a rescue amplitude *ζ*_res_(*G*) that turns its selection coefficient against the wildtype negative,

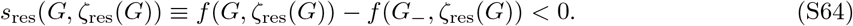

This amplitude has to be applied over a time of order of the expected time to loss of the mutant,

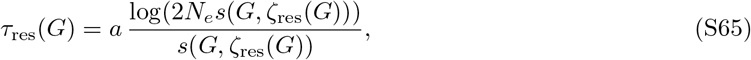

where the constant *a* is related to the expected failure rate of the rescue protocol. Together, the adaptive control protocol (*ζ*_base_, *ζ*_res_, *τ*_res_) has the time-averaged control amplitude

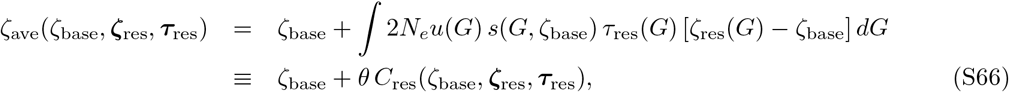

where *θ* ≡ 2*μN*_*e*_ denotes the neutral sequence diversity of the pathogen. The first term on the r.h.s. is the baseline amplitude, the second term the time-averaged extra amplitude of rescue boosts. By Eqs. (S41) and (S61), *ζ*_ave_ is linearly related to the pathogen fitness, *f_h_*, and the control efficiency, *η*. Hence, Eq. (S66) translates into a corresponding decomposition

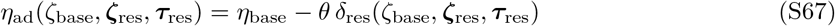

with δ_res_(*ζ*_base_, *ζ*_res_, *τ*_res_) = *c*_*p*_ *C*_res_(*ζ*_base_, *ζ*_res_, *τ*_res_)/(*q*_*ph*_*q*_*hp*_), which displays the dependence of the efficiency on the trait sequence diversity *θ*. This shows an important general point: any adaptive protocol (*ζ*_base_, *ζ*_+_, *τ*_+_) with a baseline control cost smaller in magnitude than the equilibrium host fitness, i.e., with 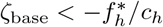, outperforms equilibrium control up to a maximum trait sequence diversity *θ*_*c*_, which is given by

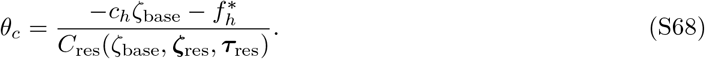

Minimizing the functional (S66) determines the optimal adaptive protocol within this class,

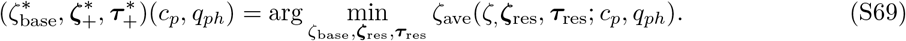

This protocol balances the baseline control cost, which is an increasing function of *ζ*_base_, and the cost of rescue boosts, which depends predominantly on the escape rates *v*_esc_ that are decreasing functions of *ζ*_base_. Evaluating optimal parameters of adaptive control becomes considerably simpler in a strong-selection regime, which generically occurs in sufficiently large pathogen populations. In this case, escape mutants are peaked at (usually few) escape channels *G*_esc_ that correspond to local maxima of *v*_esc_ and are often close to local maxima of *f_p_*. The corresponding establishment rates can be estimated from equation (S63) or be determined experimentally, for example, by Luria-Delbrück assays.

In the main text, we discuss a solvable example of adaptive ecological control in the host-pathogen fitness-model (11), (12). In this example, we fix the pathogen escape dynamics by two assumptions. (i) The mutation spectrum depends exponentially on the trait,

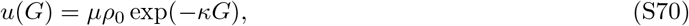

where *ρ*_0_ is an overall mutational target size. (ii) Selection is strong enough so that observed (i.e., established) escape mutants are peaked at the escape channel *G*_esc_(*ζ*_base_) = *G_e_*(*ζ*_base_), i.e., at the fitness maximum of the pathogen states with evolved resistance. Equation (S37) then determines trait effect and selection coefficient of this escape channel,

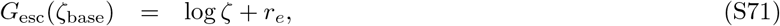

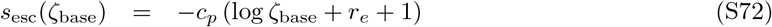

with the trait shift *r_e_* given by equation (S33). Furthermore, we consider a simplified class of protocols with a single independent parameter *ζ*_base_. In this class, the boost amplitude is set to be proportional to the baseline amplitude,

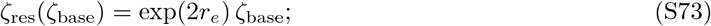

that is, the trait value *G*_esc_(*ζ*_base_) becomes the minimum of the pathogen fitness landscape *f_p_*(*G*; *ζ*_res_). The condition (S73) also fixes selection and time span of the rescue boosts as functions of *ζ*_base_,

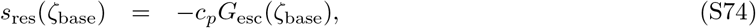

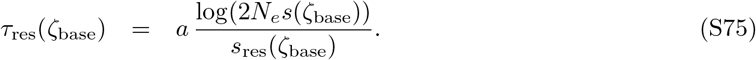

The optimization (S69) then reduces to a condition for the independent parameter *ζ*_base_,

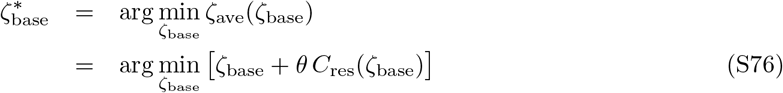

with

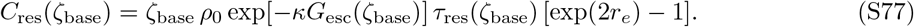

We note that, even in this simplified class of protocols, the optimization (S76) is quite complex. The baseline amplitude *ζ*_base_ tunes the effect *G*_esc_, the mutation rate *u*(*G*_esc_), and the selection coefficient *s*_esc_ of likely escape mutants, the optimal rescue of which also depends on *G*_esc_. The efficiency of the optimized adaptive protocol, 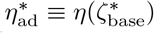, is determined by Eqs. (S41) and (S61). In Fig. 4D, we plot 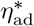 as a function of the pathogen sequence diversity *θ*, together with the efficiency *η*\* of equilibrium control (using parameters *ρ*_0_ = 1, *κ* = 1). This shows that adaptive control outperforms equilibrium control for

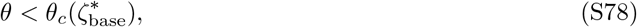

where the threshold value is determined by Eq. (S68). That is, the adaptive protocol strengthens control of pathogen populations with sufficiently small population size or mutation rate.

